# scSVA: an interactive tool for big data visualization and exploration in single-cell omics

**DOI:** 10.1101/512582

**Authors:** Marcin Tabaka, Joshua Gould, Aviv Regev

## Abstract

We present scSVA (single-cell Scalable Visualization and Analytics), a lightweight R package for interactive two- and three-dimensional visualization and exploration of massive single-cell omics data. Building in part of methods originally developed for astronomy datasets, scSVA is memory efficient for more than hundreds of millions of cells, can be run locally or in a cloud, and generates high-quality figures. In particular, we introduce a numerically efficient method for single-cell data embedding in 3D which combines an optimized implementation of diffusion maps with a 3D force-directed layout, enabling generation of 3D data visualizations at the scale of a million cells. To facilitate reproducible research, scSVA supports interactive analytics in a cloud with containerized tools. scSVA is available online at https://github.com/klarman-cell-observatory/scSVA.

---

The recent progress in the development of high-throughput single-cell methods allows researchers to study cell types and states of millions of cells, and is expected to further grow dramatically to hundreds of millions and more cells due to technological advances and the progress of focused initiatives, including the Human Cell Atlas [*1*]. Deriving biological insights from such massive datasets requires a combination of automated analysis algorithms with tools for data visualization and exploration, where users can gracefully, quickly, and iteratively explore analysis results. While substantial progress has been made on scaled analysis [*2*], far less has been done for data exploration. Indeed, existing tools [*2–4*] require extensive memory resources to explore and visualize cells and cell features, and are not scaled to massive data volumes. Here, we introduce scSVA, a lightweight R package built with Shiny for interactive visualization and exploration of single-cell omics data at the scale of up to a billion cells.

scSVA addresses common tasks in single-cell data visualization and exploration (**Fig. 1a**) by (**1**) visualizing cells on two- or three-dimensional embeddings; (**2**) visualizing gene expression values and gene signature scores on those embeddings, including by specifying a statistic to plot in a dense region with overlapping cells, such as mean or maximum gene expression value (suitable for highly or lowly expressed genes, respectively); (**3**) marking cells by clusters; (**4**) computing cell proportions and gene expression statistics for custom defined subsets of cells by polygonal selection or provided categorical data; and (**5**) producing high-quality figures combined with their comprehensive customization and annotation.

**Figure 1:**
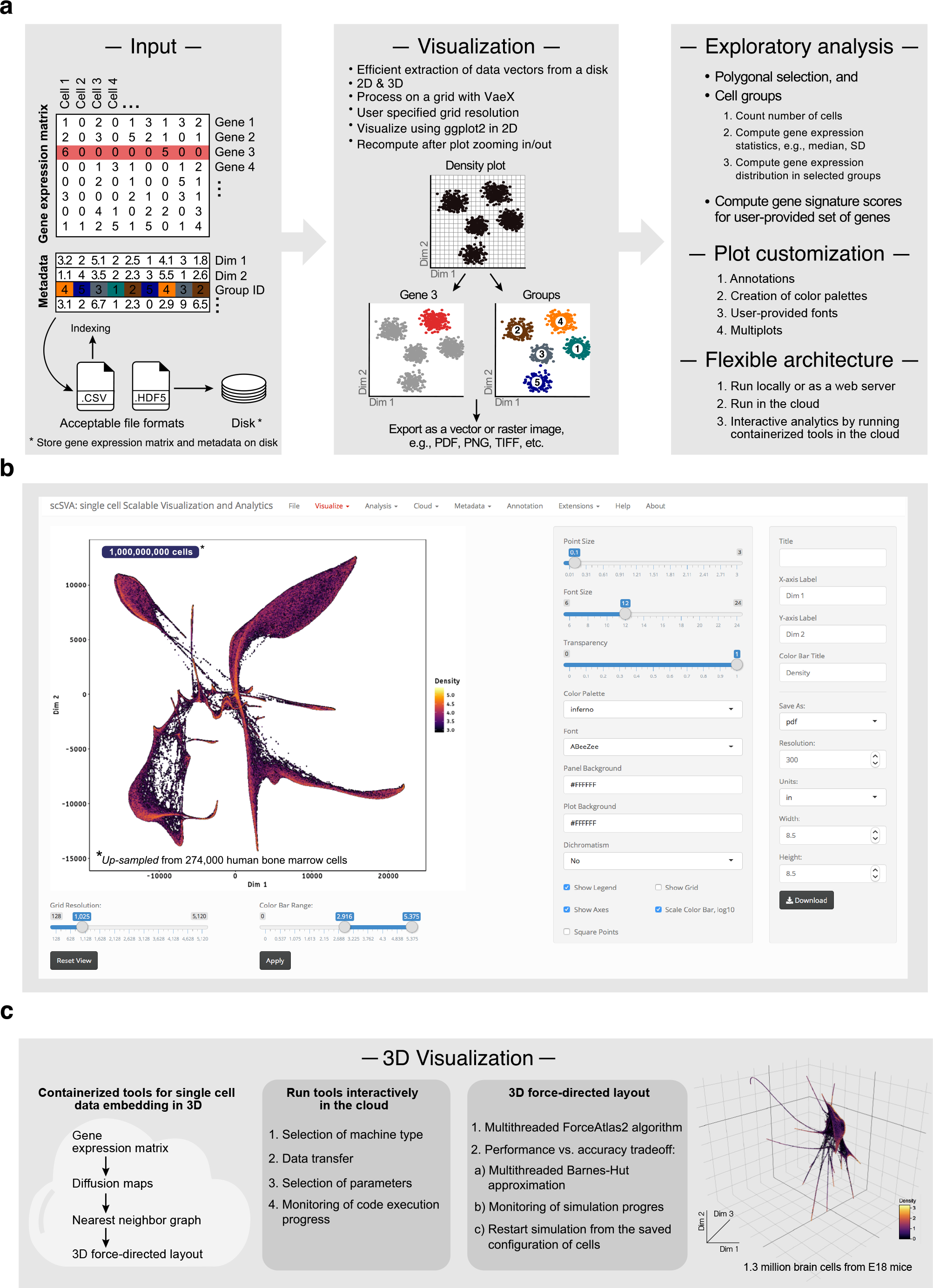
scSVA is a tool for interactive visualization and exploratory analysis of big data in single-cell omics. (**a**) Overview. scSVA accepts as input (left) an expression matrix and cell metadata in compressed text and hdf5 file formats. From disk stored datasets, scSVA retrieves only one-dimensional arrays that are needed for visualizing and exploring cell features of interest. Cells are visualized on a grid (middle) significantly reducing the number of points to be plotted. scSVA provides a set of interactive tools (right) for the exploratory analyses and plot customizations. (**b**) Interface. A screenshot of the main visualization window of the scSVA interactive application, showing 1 <u>billion</u> cell profiles, *up-sampled* for demonstration purposes from 274,000 human bone marrow cells. (**c**) 3D embeddings. Containerized code for 3D embedding can be run on a cloud platform, allowing for visualization of cells at a scale of <u>million</u> cells in 3D.

scSVA relies on advances from diverse data-heavy areas, especially astronomy, such that most of its capabilities are scaled to up to a billion cells with real time interactivity. To process cell feature vectors, scSVA uses as a back-end VaeX [*5*], an optimized python library for data processing on a grid, developed for visualization of stars brighter than magnitude 20 catalogued by the ESA space astrometry mission Gaia [*6*]. Visualizing cells on a grid allows us to effectively handle dense regions of overlapping cells on a 2D or 3D embedding and gene expression “dropouts”, common in single cell ‘omics data, especially when a user specifies a statistic to plot, such as mean or maximum gene expression level for overlapping cells. Grid data with user-specified grid resolution is visualized using R ggplot2 package (https://ggplot2.tidyverse.org), which produces aesthetically-pleasing high-quality figures. scSVA provides an extensive support for color palettes (pre-set and custom) and helps to select colors that are suitable for the color-blind. scSVA comes as a flexible and interactive Shiny application (**Fig. 1b**) enabling zooming in and out cells on a 2D embedding with immediate refreshing of cell positions on the user-defined grid resolution. We also optimized a new method for computing a single-cell data embedding in 3D which combines a new efficient implementation of diffusion maps (by optimizing the original algorithm [*7*]) with a 3D force-directed layout (ForceAtlas2 algorithm [*8*]), enabling generation of 3D data visualizations at the scale of a million cells (**Fig. 1c**).

To address computational demands, scSVA has efficient memory usage and supports interactive analytics in a cloud. To reduce memory usage, scSVA supports efficient retrieval of cell features from massive expression matrices stored on a disk (locally or on a cloud bucket) from compressed text (using indexing [*9*]) or HDF5 file formats. This reduces the memory usage by a factor equal to the number of cell features, for example ~20,000 genes in mammalian single-cell RNA-Seq gene expression matrices. For interactive analytics in a cloud (currently with Google Cloud Platform), containerized code of the tool ensures reproducible research [*10*] and eliminates the need for controlling dependencies between installed packages by target users. Thus, scSVA should enable users to interact with large datasets and complex analytics to yield novel insights and discoveries.

The scSVA package with installation instructions and extensive documentation is available online at (https://github.com/klarman-cell-observatory/scSVA) and can be installed from a source or run as a Docker container containing all system and software dependencies. Source of single-cell RNA-Seq datasets visualized in this study: the Immune Cell Atlas dataset of bone marrow immune cells (https://preview.data.humancellatlas.org) and brain cells from E18 mice (https://support.10xgenomics.com/single-cell-gene-expression/datasets).

## Competing interests

AR is a founder of, consultant to, and equity holder in Celsius Therapeutics and a member of the SAB of Syros Pharmaceuticals and ThermoFisher Scientific.

## Author Contributions

M.T. and A.R. conceived and designed the software. M.T. developed the software. M.T., J.G. and A.R conceived the 3D data visualization of single-cell datasets. M.T. and J.G. developed 3D data visualization. A.R. supervised the project.

## Acknowledgments

We thank Ayshwarya Subramanian, Christoph Muus, and Bo Li for discussions; Leslie Gaffney and Anna Hupałowska for comments and help with figures. M.T., J.G, and A.R. are supported by Klarman Cell Observatory at Broad Institute and an NHGRI CEGS grant. A.R. is an HHMI Investigator.

